# Exploring the diversity of the CO_2_-concentrating mechanism (CCM) in different C_4_ subtypes

**DOI:** 10.1101/2025.07.15.664927

**Authors:** Chiara Baccolini, Hirofumi Ishihara, Regina Feil, Leonardo Perez de Souza, Saleh Alseekh, Alisdair R. Fernie, Mark Stitt, John E. Lunn, Stéphanie Arrivault

## Abstract

C_4_ plants have traditionally been classified into NADP-malic enzyme (NADP-ME), NAD-malic enzyme (NAD-ME) and PEP carboxykinase (PEPCK) subtypes based on the predominant C_4_-acid decarboxylating enzyme. To investigate the relative contributions of malate and aspartate to C_4_-pathway fluxes in each subtype, we performed ^13^CO_2_ pulse and pulse-chase labelling experiments on four C_4_ grass species: *Zea mays* and *Setaria viridis* (NADP-ME), *Panicum miliaceum* (NAD-ME) and *Megathyrsus maximus* (PEPCK). Only a proportion (8-50%) of the total malate pool in the leaves is photosynthetically active whereas essentially all of the aspartate pool is photosynthetically active. Estimates of metabolic fluxes indicate that approximately two thirds of the C_4_ pathway flux is via malate in *Z. mays* and the remaining third via aspartate, while in *S. viridis* 50% of the flux is via malate and 50% via aspartate. In *P. miliaceum* and *M. maximus*, 91% and 85% of the flux is via aspartate and the remaining 5% and 15% via malate, respectively. The results reveal greater complexity of C_4_ pathway fluxes than is usually represented in textbook diagrams, and demonstrate the feasibility of using non-radioactive ^13^CO_2_ in pulse-chase labelling experiments to study C_4_ photosynthesis and to detect C_4_ pathway fluxes in C_3_ plants engineered to perform C_4_ photosynthesis.

**Highlight Statement:** Photosynthetic fluxes in C_4_ species are more complex than most textbook models show, with malate and aspartate both carrying C_4_ cycle fluxes in all three subtypes (NADP-ME, NAD-ME and PEPCK).

## Introduction

C4 photosynthesis is a highly efficient CO2 assimilation pathway which emerged 25-30 million years ago in the Oligocene under falling atmospheric CO2 and rising O2 concentration (Christin *et al*., 2008; Sage *et al*., 2011). Although C4 species represent only ∼3% of terrestrial plant species, they account for 23% of global primary production (Kellogg, 2013; Sage, 2016). Their high photosynthetic efficiency is driven by a biochemical CO2-concentrating mechanism (CCM) that raises the concentration of CO2 at the active site of Rubisco, thereby suppressing the oxygenation reaction and the need for photorespiration. C4 species have a specialized leaf anatomy, named Kranz anatomy, characterized by two distinct cell types: the mesophyll cells (MCs), where atmospheric CO₂ is initially fixed into C_4_ acids (malate and aspartate), and the bundle sheath cells (BSCs), where CO₂ is released by C_4_ acid decarboxylation and re-fixed by Rubisco (Hatch and Osmond, 1976). The C_4_ CCM brings several advantages including faster photosynthesis, lower photorespiration, and increased water and nitrogen use efficiencies (von Caemmerer and Furbank, 2003). Due to these advantages, engineering C_4_ photosynthesis into C_3_ crops is considered a promising approach to boost crop productivity and improve resilience in the face of global climate change (Covshoff and Hibberd, 2012; Cui, 2021; Furbank *et al*., 2023).

In C_4_ plants, atmospheric CO2 is converted to bicarbonate (HCO ^-^) by carbonic anhydrase in the MC, which is then assimilated by phospho*enol*pyruvate carboxylase (PEPC) to produce oxaloacetate (OAA), a C_4_ acid. Oxaloacetate is converted to malate or aspartate, which diffuse via plasmodesmata into the BSC, where they are decarboxylated to release CO_2_, which is then re-fixed by Rubisco. The residual C3 moiety diffuses back to the MC to regenerate the initial CO2 acceptor, PEP. While all C4 plants employ PEPC for initial CO2 fixation into C_4_ acids, C_4_ species differ in the mode of C_4_ acid decarboxylation and have conventionally been divided into three subtypes according to the predominant decarboxylating enzyme (Hatch, 1987): NADP malic enzyme (NADP-ME), NAD-malic enzyme (NAD-ME) or phospho*enol*pyruvate carboxykinase (PEPCK) (Figure 1A). Linked to the mode of decarboxylation, there are also differences in the inter- and sub-cellular compartmentation of C_4_ pathway reactions and which C_4_ and C_3_ metabolites move between the cells. This metabolic diversity reflects the independent evolution of C4 photosynthesis in over 60 separate plant lineages (Sage, 2016).

**Figure 1.**
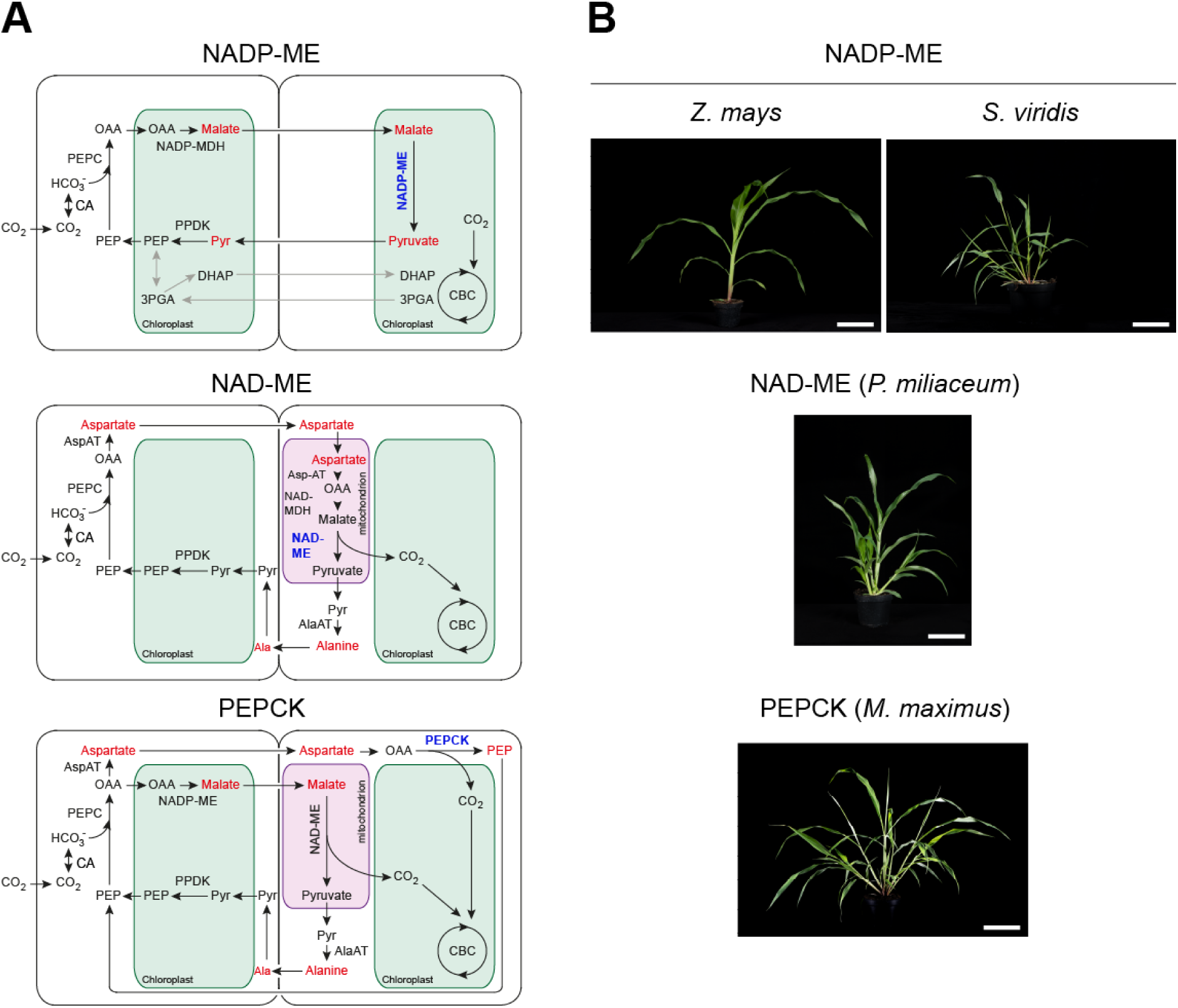
Photosynthetic pathways in the three C_4_ subtypes. (**A**) Inter- and subcellular compartmentation of the C_4_ pathways in canonical NADP-malic enzyme (NADP-ME), NAD-malic enzyme (NAD-ME) and phosphoenolpyruvate carboxykinase (PEPCK)-type C_4_ species. Abbreviations: 3PGA, 3-phosphoglycerate; Ala, alanine; Ala-AT, alanine aminotransferase; Asp-AT, aspartate aminotransferase; CA, carbonic anhydrase; CBC, Calvin-Benson cycle; DHAP, dihydroxyacetone-phosphate; NADP-MDH, NADP-dependent malate dehydrogenase; OAA, oxaloacetate; PEP, phospho*enol*pyruvate; PEPC, PEP carboxylase; PPDK, pyruvate, phosphate dikinase; Pyr, pyruvate. The predominant C_4_ acid and C_3_ products of decarboxylation are shown in red and the major decarboxylating enzymes in blue. The 3PGA:triose-phosphate energy shuttle in NADP-ME species is shown by the grey arrows. (**B**) Photographs of *Zea mays* and *Setaria viridis* (NADP-ME subtype), *Panicum miliaceum* (NAD-ME subtype) and *Megathyrsus maximus* (PEPCK subtype) plants used in this study (bar = 10 cm).

In the NADP-ME subtype, textbook diagrams generally show OAA being reduced to malate by NADP-malate dehydrogenase (NADP-MDH) in the MC chloroplasts, and malate being the sole C4 acid that diffuses into the BSC for decarboxylation by NADP-ME in the BSC chloroplasts, releasing CO_2_, NADPH and pyruvate (Figure 1A). Pyruvate returns to the MC to regenerate PEP via pyruvate, phosphate dikinase (PPDK) in the MC chloroplasts. In *Z. mays* and some other NADP-ME species, there is insufficient NADPH production in the BSC chloroplasts to convert all of the 3-phosphoglycerate (3PGA) into triose-phosphates, so some of the 3PGA moves to the MC chloroplasts for reduction to triose-phosphates, which then move back to the BSCs to re-enter the Calvin-Benson cycle (CBC) (Leegood, 1985; Wang *et al*., 2014; Majeran *et al*., 2015). In the NAD-ME subtype, textbook diagrams show OAA being transaminated in the MC cytosol by aspartate aminotransferase (AspAT) to produce aspartate, which diffuses into the BSC (Figure 1A). There, it is transported into the mitochondria, where it is first converted back to OAA by AspAT, then reduced to malate by NAD-malate dehydrogenase (NAD-MDH), and finally decarboxylated by NAD-ME to release CO_2_, NADH and pyruvate. The pyruvate is converted to alanine by alanine aminotransferase (AlaAT) which returns to the MC, where it is deaminated by AlaAT to pyruvate that is used by PPDK for PEP regeneration. The C_4_ pathway is more complicated in the PEPCK subtype, where both malate and aspartate are produced from OAA and diffuse into the BSCs (Figure 1A). Aspartate diffuses to the BSC cytosol, where it is deaminated by AspAT and decarboxylated by PEPCK in the cytosol, releasing CO2 and PEP. Textbook diagrams typically show PEP moving back to the MC to be used directly by PEPC, but the form in which the residual C_3_ moiety moves back to the MCs in PEPCK species is uncertain. In most PEPCK subtypes, malate moves to the BSC mitochondria where is it decarboxylated by NAD-ME, releasing CO2, NADH and pyruvate. Pyruvate is converted by AlaAT to alanine that moves to the MC, where it is deaminated to pyruvate, which is used to regenerate PEP. In some PEPCK-type species, malate is decarboxylated by NADP-ME in the BSC chloroplasts. The NAD-ME and the PEPCK subtypes require intercellular amino acid shuttles to maintain nitrogen balance. AspAT and AlaAT use glutamate and 2-oxoglutarate (2OG) as amino donor and acceptor, respectively.

The C_4_ pathway was discovered by ^14^CO_2_ labelling experiments in sugarcane (*Saccharum officinarum*), an NADP-ME species (Kortschak *et al*., 1965; Hatch and Slack, 1966; Johnson and Hatch, 1969). The identification of some other species as C_4_ plants was also based on early ^14^CO_2_ pulse-labelling studies showing initial fixation of CO_2_ into C_4_ acids rather than 3PGA. However, elucidation of the NAD-ME- and PEPCK-type C_4_ pathways and assignment of C_4_ species to a given subtype were based primarily on measurements of maximal extractable enzyme activities and metabolite feeding experiments with isolated BSC strands (Hatch *et al*., 1975; Furbank *et al*., 1990; Carnal *et al*., 1993), supported in only a few species, such as *Spartina* spp. (a putative PEPCK species) (Thomas and Long, 1978; Smith *et al*., 1982) by ^14^CO_2_ labelling experiments on intact leaves. Some previous labelling studies revealed a more complex picture of C_4_ pathway fluxes than expected from the textbook models. Based on ^14^CO_2_ pulse-chase labelling studies, Hatch (1971) reported substantial movement and decarboxylation of aspartate as well as malate in *Z. mays*, while Meister *et al*. (1996) showed that 35-40% of the C_4_ pathway flux in another model NADP-ME species, *Flaveria bidentis*, is carried via aspartate. A recent ^13^CO_2_ pulse-labelling study in *Z. mays* indicated that the contribution of aspartate to overall C_4_ pathway flux ranges from 14 to 44% (Weissman *et al*., 2016; Arrivault *et al*., 2017; Medeiros *et al*., 2022). It was further shown that the relative contributions of malate and aspartate to C_4_ pathway flux in *Z. may*s is dependent on the environmental conditions, with aspartate becoming the predominant C_4_ acid under low irradiance (Medeiros *et al*., 2022). These findings, showing unexpected diversity in the C_4_ pathways in NADP-ME-type species, support an emerging view that the conventional division of C_4_ species into three subtypes is an over-simplification (Furbank, 2011), a view reinforced by transcript profiling and metabolic modelling of C_4_ species (Pick *et al*., 2011; Wang *et al*., 2014). However, to date, there has been little direct examination of C_4_ pathway fluxes in NAD-ME and PEPCK species, to assess whether these fit with the standard textbook models.

The aim of this study was to determine the relative contributions of malate and aspartate to C_4_ pathway fluxes in representative species from the three subtypes, using ^13^CO_2_ pulse and pulse-chase labelling approaches. To minimize phylogenetic differences, we selected four closely related species from the Panicoideae subfamily within the Poaceae (grasses): *Z. mays* (NADP-ME), *Setaria viridis* (green foxtail millet; NADP-ME), *Panicum miliaceum* (proso millet; NAD-ME) and *Megathyrsus maximus* (guinea grass, previously known as *Panicum maximum*; PEPCK) (Figure 1B). *Z. mays* has been widely used as a model species for studies of C_4_ photosynthesis, and protocols for ^13^CO_2_ pulse-labelling experiments have been established in this species (Medeiros *et al*., 2022). There is considerable interest in introducing the C_4_ pathway into C_3_ crop species, such as rice, to improve their photosynthetic efficiency and productivity (Furbank *et al*., 2023), with the *Z. mays* NADP-ME pathway providing the preferred blueprint (Lin *et al*., 2020; Ermakova *et al*., 2021). Therefore, it is important to have a fuller understanding of how the C_4_ pathway actually operates in this species. *S. viridis* has been adopted as a general model for genetic studies in C_4_ plants, due to its short life-cycle and relative ease of transformation (Brutnell *et al*., 2010). Although classified as an NADP-ME species, surprisingly little is known about its C_4_ photosynthetic pathway. The textbook pathways for NAD-ME and PEPCK subtypes are based to a large extent on *in-vitro* experiments with *P. miliaceum* and *M. maximus* BSC strands, respectively, and these two species are commonly used as representatives of these two subtypes (Furbank *et al*. 1990; Carnal *et al*., 1993).

Our ^13^C labelling data show that C_4_ pathway fluxes are more complex than usually described in textbook models, with parallel malate and aspartate pathways occurring in all three subtypes, even in the canonical NADP-ME species *Z. mays*. Our results also validate the use of ^13^CO_2_ as an alternative to radioactive ^14^CO_2_ for pulse and pulse-chase labelling approaches to investigate C_4_ pathway fluxes in C_4_ species, and also to detect C_4_ fluxes in transgenic C_3_ plants that have been engineered to perform C_4_ photosynthesis.

## Materials and methods

### Plant materials and chemicals

Seeds were obtained from the following sources: *Zea mays* L. cv B73 from KWS SAAT SE & Co. KGaA (Einbeck, Germany); *Setaria viridis* (L.) P.Beauv. accession A10.1 from Prof. Steve Kelly (University of Oxford, UK); *Panicum miliaceum* L. from Spreewälder BioMühle GmbH iG (Kolkwitz, Germany); and *Megathyrsus maximus* (Jacq.) B.K.Simon & S.W.L.Jacobs accession 78380 from the Royal Botanic Gardens, Kew (www.kew.org). Gas cylinders containing N_2_, O_2_ or unlabelled CO_2_ (^12^CO_2_) were obtained from Air Liquide Deutschland GmbH (Düsseldorf, Germany; https://de.airliquide.com), while ^13^CO_2_ (isotopic purity 99 atom %) was obtained from Sigma-Aldrich (www.sigmaaldrich.com). Enzymes and biochemicals were obtained from Sigma-Aldrich, Roche Applied Science (lifescience.roche.com) or Merck (www.merckmillipore.com).

### Plant growth

1. *Z. mays* seeds were germinated in darkness in petri dishes on moistened filter paper (3 days, 28° C). The imbibed seeds were sown into a soil mixture comprised of natural clay, white peat and wood fibres (Einheitserde Typ Topf; www.einheitserde.de) mixed 2:1 with quartz sand, with one seed per 10-cm diameter pot, and germinated under long-day conditions (16-h photoperiod) with an irradiance of 105 µE m^-2^ s^-1^, day/night temperatures of 22° C/18° C and 70% relative humidity). After 5 days, the seedlings were transferred to 14 h/10 h day/night cycles with an irradiance of 550 µE m^-2^ s^-1^ provided by white LEDs tuned to a sunlight-like spectrum, supplemented with 60 µE m^-2^ s^-1^ of far-red light, with day/night temperatures of 28° C/20° C, and 80% relative humidity. The fourth fully expanded leaves of three-week-old plants were used for ^13^CO_2_ labelling.
2. *S. viridis* seeds were incubated in moist sphagnum moss at 4° C for 3 weeks to promote germination. The pre-treated seeds were cleaned, dried overnight at 37° C, and germinated on wet tissue paper under 14 h/10 h day/night cycles (irradiance 550 µE m^-2^ s^-1^ and 60 µE m^-2^ s^-1^ far-red light, 28° C/20° C, 80% relative humidity). After 2-3 days, the germinated seedlings were transferred to soil (as described above for *Z. mays*) in 10-cm diameter pots, with one seedling per pot. The plants were grown under the same conditions as during germination until 2 weeks old, when they were used for ^13^CO_2_ labelling experiments.
3. *P. miliaceum* and *M. maximus* seeds were germinated in petri dishes on moistened filtered paper under 13-h or 16-h photoperiod, respectively, with an irradiance of 40 µE m^-2^ s^-1^ (white LEDs) plus 60 µE m^-2^ s^-1^ far-red light, day/night temperatures of 24° C/20° C, and 60% relative humidity. After ten days, the seedlings were transferred to soil (as described above for *Z. mays*) and grown under a 13-h photoperiod with an irradiance of 550 µE m^-2^ s^-1^ plus 60 µE m^-2^ s^-1^ far-red light, day/night temperatures of 24° C/20° C and 60% relative humidity. The fourth fully expanded leaf of 3-4 week old plants was used for ^13^CO_2_ labelling experiments. The four species were grown at separate times in the same growth chamber under identical irradiance conditions, but with slightly different photoperiods to delay flowering.

### ^13^CO_2_ pulse, ^13^CO_2_ pulse-^12^CO_2_ chase labelling experiments

Labelling experiments were performed *in situ* in the growth chamber as described in Baccolini and Arrivault (2024). Labelling started a minimum of 2 h after the start of the light period to ensure that photosynthetic rates were at steady state. For labelling of *Z. mays* leaves, the mid-section of the fourth fully expanded leaf was clamped in a horizontal orientation in a custom-built Perspex labelling chamber (Ermakova *et al*., 2021) while still attached to the mother plant. For the other three species, which have narrower leaves, two to three leaves were used. The leaves received an irradiance of 510 µE m^-2^ s^-1^ inside the labelling chamber, and were flushed with a humidified artificial air mixture containing 79% N_2_, 21% O_2_ and ^12^CO_2_ (420 µmol mol^-1^) for 60 s at a flow rate of 10 L min^-1^ to equilibrate. For pulse-labelling, the gas supply inside the chamber was switched to a humidified air mixture containing 79% N_2_, 21% O_2_ and ^13^CO_2_ (420 µmol mol^-1^) at a flow rate of 10 L min^-1^. Instantaneous switching between labelled and unlabelled air flows was achieved using a 4-way valve Ermakova *et al*., 2021). After the required pulse-labelling time, the leaves were rapidly quenched by pouring liquid N_2_ into the labelling chamber through an opening in the top plate, via a funnel fitted with a removable plug, with a liquid N_2_ exit hole located on the opposite side of the lower plate to ensure flow of liquid N_2_ through the whole chamber. The frozen leaf was cut from the mother plant, trimmed to remove the basal and distal parts that had been covered by the seals at the edge of the labelling chamber, and then stored at −80° C until analysis. The pulse labelling times for *S. viridis* were: 5, 10, 20, 30 s, and 1, 3, 5, 10, 20, 40 or 60 min, while those for *P. miliaceum* and *M. maximus* were: 5, 10, 15, 20, 30, 50 s, and 1, 3, 5, 10, 20, 40 or 60 min. For pulse-chase labelling, the leaves were exposed to air mixtures containing ^13^CO_2_ for 20 s, as described above, and then the gas supply was switched to an unlabelled air mixture containing 420 µmol mol^-1^ ^12^CO_2_ for the chase. After 15, 30, 60. 90 or 120 s chase, the leaves were rapidly quenched with liquid N_2_, and processed as above. Unlabelled control samples (t_0_) were collected for all species by equilibrating leaves in the labelling chamber for 60 s, as described above, and then quenching with liquid N_2_. The data from ^13^CO_2_ pulse-labelling of *Z. mays* are from the experiments described in Medeiros *et al*. (2022). The *Z. mays* pulse-chase labelling was performed under identical conditions on a new batch of plants, with all of the labelling completed within one day. For *S. viridis*, *P. miliaceum* and *M. maximus*, the pulse- and pulse-chase labelling experiments were performed on a single batch of plants of each species, with the different pulse- and pulse-chase labelling times being randomized over two consecutive days. A fresh plant was used for each labelling treatment, and at least three biological replicates were collected for each treatment.

### Metabolite analysis

The frozen leaf tissue was ground to fine powder using a ball mill (Tesch, Haan, Germany) at liquid N_2_ temperature. For mass spectrometry analysis, aliquots (15-20 mg) of frozen tissue powder were extracted with chloroform-methanol as described in Arrivault *et al*. (2009). Isotopologues of phosphorylated intermediates, malate, pyruvate, 2-oxoglutarate and aspartate were quantified by anion-exchange high-performance liquid chromatography coupled to tandem mass spectrometry (LC-MS/MS) on an AB Sciex 6500 Q-Trap triple quadrupole mass spectrometer as described in Lunn *et al*. (2006) with modifications as described in Figueroa *et al*. (2016) or by reverse-phase ion-pair LC-MS/MS as described in Arrivault *et al*. (2017). Isotopologues of alanine and glutamate were quantified by gas chromatography coupled to mass spectrometry as described in Lisec *et al*. (2006). The total amounts of PEP, pyruvate and 3PGA were also determined enzymatically in freshly prepared trichloroacetic acid extracts as described in Merlo *et al*. (1993) using a UV-Vis spectrophotometer (Shimadzu, Kyoto, Japan). The relative isotopologue distributions and ^13^C amounts (natom ^13^C equivalents g^-1^ FW) in each metabolite were calculated as in Arrivault *et al*. (2017).

### Chlorophyll and protein determination

Chlorophyll was extracted by adding 600 µl methanol to 20 mg FW frozen leaf tissue powder, followed by vortex mixing and incubation in the dark for 30 min at 4° C. Insoluble cell debris was pelleted by centrifugation at 14,000x*g* for 5 min (4° C). An aliquot (150 µl) of the supernatant was taken for measurement of the absorbance at 650 nm and 665 nm. The chlorophyll a and b contents were calculated using the formulae s described in Porra *et al*. (1989). Proteins were extracted from 20 mg FW frozen leaf tissue powder by addition of 750 µl extraction buffer containing: 0.1 M Tris-HCl, pH 8, 0.2 M NaCl, 5 mM EDTA, 2% (w/v) SDS, 17.5 mM *β*-mercaptoethanol and protease inhibitor cocktail (P9599, Sigma, Germany; www.sigmaaldrich.com). After vortex mixing, the suspension was incubated for 30 min at room temperature, re-mixed, and then the insoluble cell debris was pelleted by centrifugation at 1500×*g* for 10 min at room temperature. The supernatant was decanted into a new tube and the pellet was re-extracted twice more with 750 µL of extraction buffer. The three supernatants from the successive extractions were pooled for the protein measurement. Protein was quantified colorimetrically using a reducing-agent-compatible form of the bicinchoninic acid assay reagent (Pierce BCA Protein Assay; Thermo Fisher Scientific, Germany; www.thermofisher.com) with bovine serum albumin as standard.

### Measurement of photosynthetic CO_2_ assimilation rates

Photosynthetic CO_2_ assimilation rates were measured in the fourth fully expanded leaves of 3 to 4-week-old *S. viridis*, *P. miliaceum* and *M. maximus* plants using an LI-6400XT open-flow infrared gas exchange analyser system (LI-COR Inc., Lincoln, NE, USA; www.licor.com) equipped with a 2-cm^3^ LI-6400-40 leaf chamber fluorometer (LI-COR Inc.).Light response curves (in a range from 0-2000 µE m^-2^ s^-1^) were measured at a constant CO_2_ concentration of 420 µmol mol^-1^, with a relative humidity at 50% and a leaf temperature of 26° C. The amount of blue light was set to 10% photosynthetic photon flux density to ensure stomatal opening.

### Statistical analyses

Statistical analyses were performed using Prism 10 software (GraphPad Software, Boston, MA, USA; https://www.graphpad.com). Specific information on the statistical tests used for each dataset is reported in the corresponding figure legends.

## Results

### ^13^CO_2_ pulse-labelling experiments

Plants were grown in a controlled environment chamber fitted with LED lamps tuned to a sunlight-like spectrum, giving an irradiance of 500-550 µE m^-2^ s^-1^. We performed ^13^CO_2_ pulse-labelling experiments on *S. viridis*, *P. miliaceum* and *M. maximus* plants *in situ* within the growth chamber. Labelling was done on the fourth fully-expanded leaf under steady state photosynthetic conditions at an irradiance of 510 µE m^-2^ s^-1^. At this irradiance, CO_2_ assimilation rates were approximately 62%, 60% and 74% of the light-saturated rates in *S. viridis*, *P. miliaceum* and *M. maximus*, respectively (Supplementary Figure S1; Supplementary Dataset S1A), consistent with previous reports that light-saturated CO_2_ assimilation rates are generally higher in NADP-ME and NAD-ME species than in PEPCK species (Edwards *et al*., 2001). There were only minor differences between species in their leaf protein and chlorophyll contents (Supplementary Dataset S1B). Leaves were supplied with air containing ^13^CO_2_ (420 µmol mol^-1^) for various times ranging from 5 s to 60 min., then rapidly quenched *in situ* with liquid nitrogen and extracted. The isotopic composition of key metabolites from the C_4_ pathway (malate, aspartate, pyruvate, alanine, PEP, glutamate and 2OG) and CBC (3PGA and DHAP) was analysed using high-performance liquid chromatography coupled to tandem mass spectrometry (LC-MS/MS) and gas-chromatography coupled to mass spectrometry (GC-MS). Isotopologue amounts (nmol g^-1^), total metabolite amounts (nmol g^-1^), isotopologue distributions (%) and ^13^C enrichments (%) are presented in Supplementary Datasets S2A, B, C and D, respectively. Data from a ^13^CO_2_ pulse-labelling experiment on *Z. mays* plants that were grown and labelled under identical conditions to the other three species were available from a previous study (Medeiros *et al*., 2022). The published *Z. mays* labelling data were re-analysed for comparison with the new ^13^CO_2_ pulse-labelling data from the other three species.

Our LC-MS/MS analysis of isotopologues does not provide positional information. Given the very low (<1%) natural abundance of the heavy isotopes of hydrogen (^2^H) and oxygen (^17^O and ^18^O), the m_0_ (unlabelled) and m_4_ (fully labelled) isotopologues of malate (or aspartate) that we measured essentially represent single isotopomers with zero or four ^13^C atoms, respectively. In contrast, the m_1_, m_2_ and m_3_ isotopologues of malate comprise multiple isotopomers with one, two or three ^13^C atoms, respectively, distributed in different positions within the molecule. For example, the measured m_1_ isotopologue of malate could include [1-^13^C]malate, [2-^13^C]malate, [3-^13^C]malate and [4-^13^C]malate isotopomers. However, during short-term (≤ 20 s) pulse-labelling with ^13^CO_2_, only the C4 position of malate and aspartate will be directly labelled via carboxylation of PEP by PEPC. Therefore, for calculation purposes, it was a reasonable assumption that the m_1_ isotopologues of malate and aspartate comprised only the respective C4 isotopomers (i.e., [4-^13^C]malate and [4-^3^C]aspartate). Therefore, we used the m_1_ isotopologue data to estimate the initial rates of CO_2_ fixation into the C_4_ acids.

Label was rapidly incorporated into both malate and aspartate in all the C_4_ subtypes (Figure 2A), with the respective m_1_ isotopologues being the only labelled forms after short (≤20 s) pulses. These results show that both malate and aspartate are early products of primary CO_2_ fixation in all four species. In all subtypes, the abundance of aspartate m_0_ declined in a monophasic manner to a very low value at the end of the 60 min pulse, and with values below 10% (*Z. mays*, *S. viridis, P. miliaceum*) and 20% (*M. maximus*) within 120 s. This is consistent with the presence of a single metabolically active pool that is engaged in the C_4_ pathway. In contrast, malate m_0_ declined more slowly and was still falling when aspartate m_0_ had approached zero. This pattern indicates the presence of at least two malate pools, one that is photosynthetically active and rapidly labelled, and another that is not directly involved in photosynthetic C fixation and is labelled much more slowly. We will refer to these as the photosynthetically active and inactive malate pools, respectively. Labelling data for other C_4_ pathway intermediates and CBC intermediates are presented in Supplementary Figure S3.

**Figure 2.**
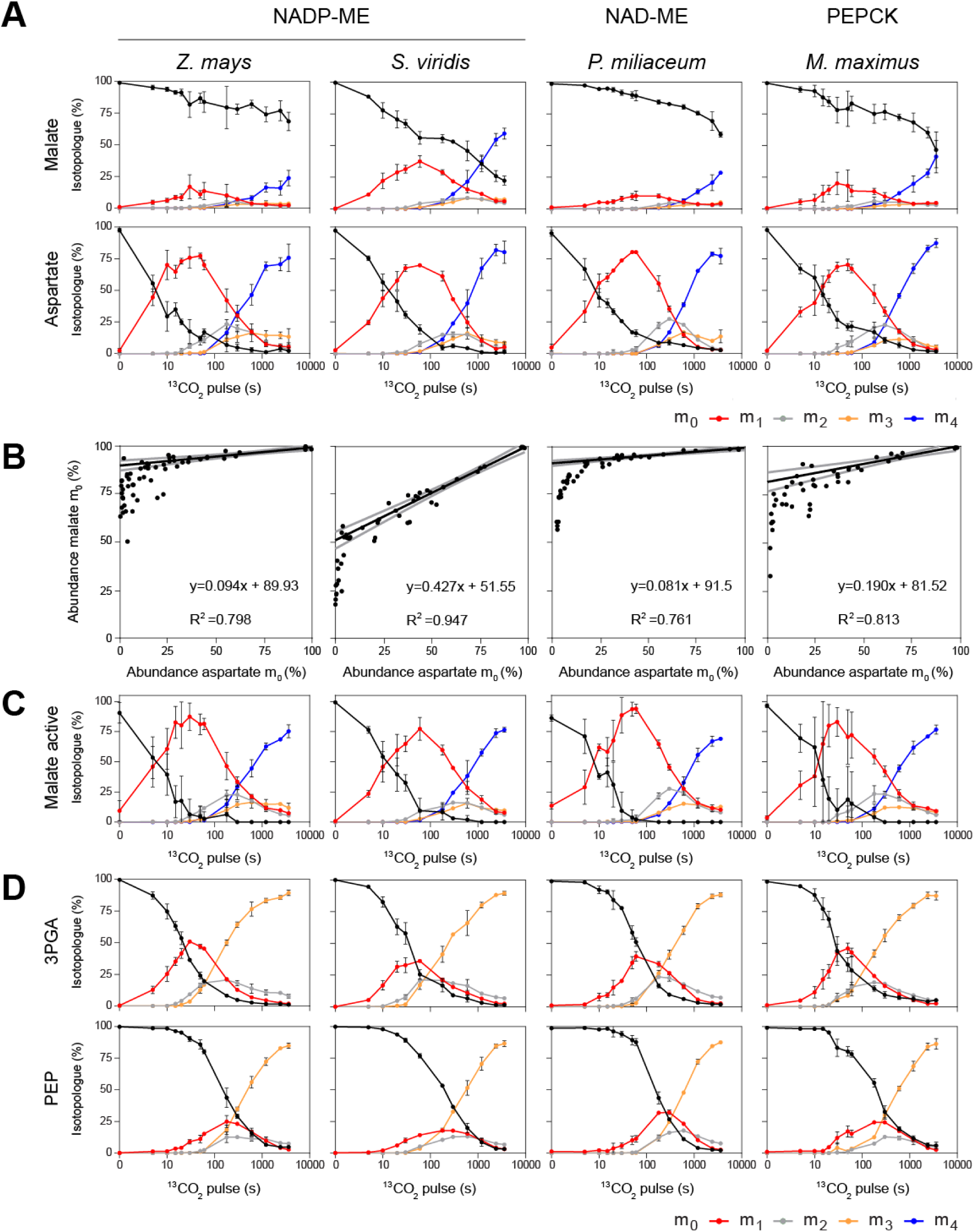
^13^CO_2_ pulse-labelling of photosynthetic intermediates in C_4_ species. *Zea mays, Setaria viridis* (NADP-ME subtype), *Panicum miliaceum* (NAD-ME subtype) and *Megathyrsus maximus* (PEPCK subtype) leaves were pulse labelled with ^13^CO_2_ under steady state photosynthetic conditions. Data for *Z. mays* are from a previous study (Medeiros *et al*., 2022). (**A**) Labelling kinetics of malate and aspartate (n.b. the time axis is shown on a log_10_ scale). (**B**) Determination of the photosynthetically active malate pool size in each species by comparing the abundance of the unlabelled m_0_ isotopologues of malate and aspartate in each sample. Lines were fitted to the data from the short-duration pulse times (0-20s) by linear regression to determine the y-axis intercept. The 95% confidence intervals are indicated by grey lines, and the equations of the fitted lines and regression coefficients (R^2^) are shown. (**C**) Labelling kinetics of malate after normalization to the photosynthetically active pool size. (**D**) Labelling kinetics of 3PGA and PEP. In (**A**), (**C**) and (**D**), the relative abundance of each isotopologue (*m_x_*) is shown as a percentage of the total abundance of the metabolite, where *x* is the number of ^13^C atoms present. Data are shown as the mean ± SD (*n*= 2 to 11). The original isotopologue abundance data are in Supplementary Dataset S2C.

### ^13^CO_2_-pulse/^12^CO_2_-chase labelling experiments

We applied a classical pulse-chase labelling approach to follow the movement of carbon through the C_4_ pathway after the initial fixation into malate and aspartate. Leaves from all four species were pulse-labelled with ^13^CO_2_ for 20 s before switching to an unlabelled air mixture containing ^12^CO_2_ (420 µmol mol^-1^) for the chase, with various chase times ranging from 10-120 s. The duration of the ^13^CO_2_ pulse was chosen to ensure that there was readily detectable labelling of malate and aspartate, but negligible labelling of the C_4_ acids in the C1-C3 positions. In all four species, we observed a rapid decrease in the abundance of the m_1_ isotopologues of malate and aspartate during the chase, accompanied by a rise in the abundance of the unlabelled m_0_ isotopologue (Figure 3A). There were no significant changes in the labelling of pyruvate or alanine during the chase (Supplementary Figure S4). The changes in the abundance of the m_1_ isotopologues indicate specific labelling of malate and aspartate in the C4 position during the pulse, followed by rapid loss of ^13^C from the C4 position during the chase, leaving the residual C_3_ moiety (pyruvate or alanine) unlabelled, as would be expected from decarboxylation of the C_4_ acids. There was a small transient rise in the abundance of the m_1_ isotopologue of 3PGA in *Z. mays* and *M. maximus* at the beginning of the chase, followed by a gradual decline, while in *S. viridis* and *P. miliaceum* the labelling of 3PGA declined after an initial lag (Figure 3B). A transient rise in labelling of another CBC intermediate, DHAP (m_1_), was also observed in all four species at the beginning of the chase (Figure 3B). These labelling patterns resemble those reported from classical ^14^CO_2_ pulse-chase labelling experiments in C_4_ plants (Johnson and Hatch, 1969), indicating re-fixation of ^13^CO_2_ by Rubisco (into 3PGA) following its release by decarboxylation of C_4_ acids in the BSCs.

**Figure 3.**
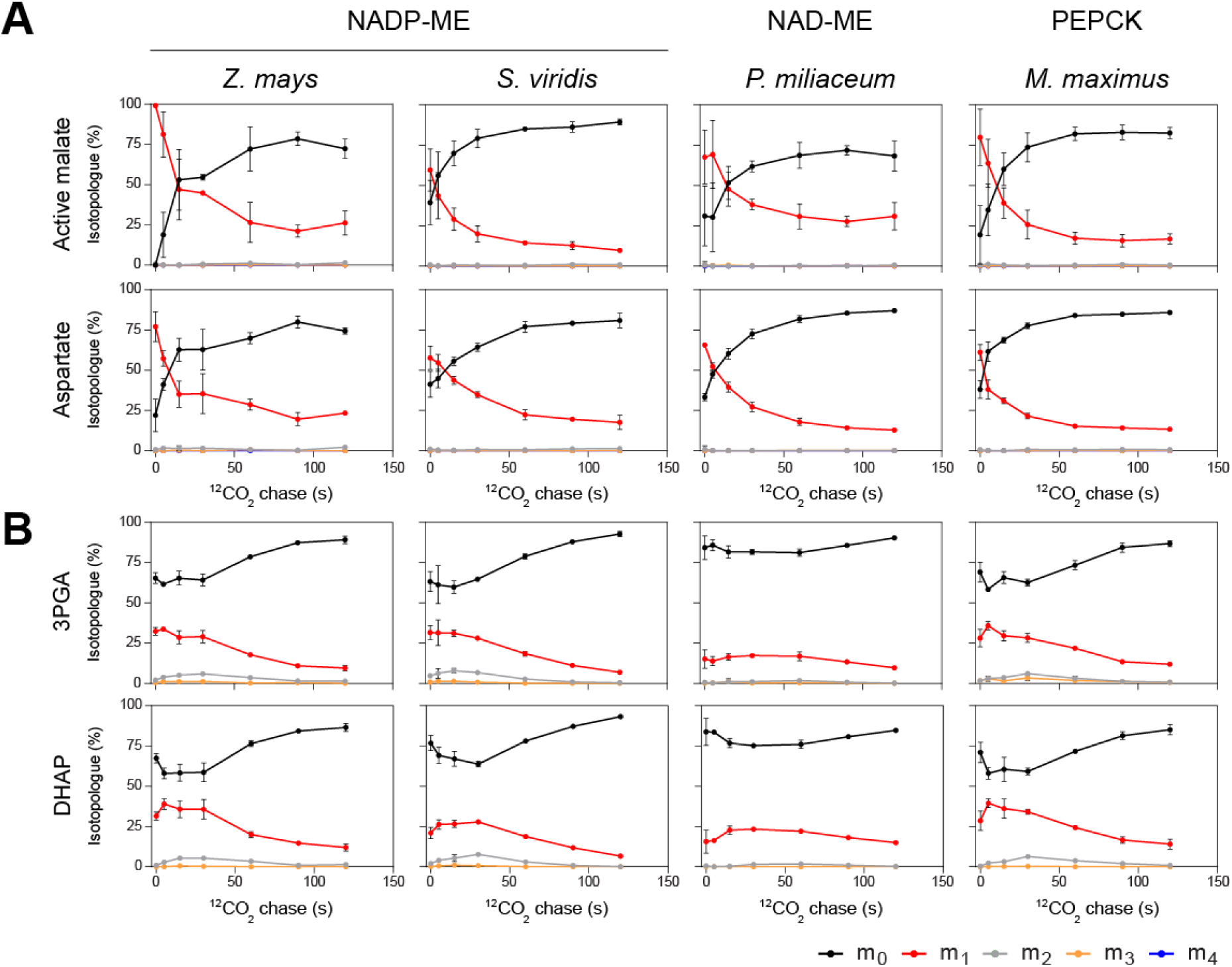
^13^CO_2_ pulse-chase labelling of photosynthetic intermediates in C_4_ species. *Zea mays, Setaria viridis* (NADP-ME subtype), *Panicum miliaceum* (NAD-ME subtype) and *Megathyrsus maximus* (PEPCK subtype) leaves were pulse labelled with ^13^CO_2_ under steady state photosynthetic conditions for 20 s, followed by a chase (5-120 s) in air containing ^12^CO_2_. (**A**) Labelling kinetics of malate and aspartate. The malate data are normalized to the photosynthetically active malate pool size, determined as described in Figure 2B. (**B**) Labelling kinetics of 3GA and DHAP. The relative abundance of each isotopologue (*m_x_*) is shown as a percentage of the total abundance of the metabolite, where *x* is the number of ^13^C atoms present. Data are shown as the mean ± SD (*n*= 2 to 11). n.b. in both panels the time axis is shown on a log_10_ scale. The original isotopologue abundance data are in Supplementary Dataset S2C.

### Estimation of the photosynthetically active malate pools by comparison with aspartate labelling kinetics

The presence of inactive pools that are not involved in photosynthesis complicates the analysis of ^13^CO_2_ labelling kinetics. This is especially the case for malate, where *Z. mays* leaves contain large pools in the vacuole and non-photosynthetic cell types (Leegood, 1985; Weiner and Heldt, 1992) and the overall pool is only partly labelled (Weissmann et al., 2016; Arrivault et al., 2017, Medeiros et al., 2022).

The size of the photosynthetically active malate pool in *Z. mays* leaves was previously estimated from changes in the abundance of the unlabelled (m_0_) isotopologue as it declined to a near-asymptotic level, but not zero, after ^13^CO_2_ pulse labelling for 30-60 min (Arrivault *et al*., 2017; Medeiros *et al*., 2022). However, this approximation could not be applied to the labelling data from *S. viridis*, *P. miliaceum* and *M. maximus,* as the malate m_0_ isotopologue abundance continued to fall throughout the pulse-labelling period in these species. Thus, an alternative approach was used to estimate the size of the photosynthetically active malate pool (Figure 2B). We plotted aspartate m_0_ abundance (x axis) against malate m_0_ abundance (y axis) and calculated, by linear regression, the slope and y-axis intercept for the samples labelled within the first 20 s (n.b. the regression coefficient (R^2^) was substantially lower if later time points were also included in the analysis). Calculating [100 minus the intercept] gives an estimate of the photosynthetically active pool of malate (expressed as a percentage of the total malate pool) that was rapidly labelled with similar kinetics to aspartate. Applying this approach to the previously published pulse-labelling data from *Z. mays*, we estimated that the photosynthetically active malate pool represented about 10% of total malate (Figure 2B). This value was somewhat lower than our previous estimations from the same datasets (based on the asymptotic value of the m_0_ isotopologue of malate), which were in a range from 24-27% (Arrivault *et al*., 2017; Medeiros *et al*., 2022). This discrepancy might be explained by the presence of more than two pools of malate: (1) a small pool of photosynthetically active malate that labels rapidly, (2) another small metabolically active pool that labels more slowly but is essentially fully labelled by 60 min, and (3) a larger, metabolically inactive pool that remains essentially unlabelled. In this scenario, using the asymptotic value of the malate m_0_ isotopologue after a 60-min pulse would lead to over-estimation of the photosynthetically active pool size because the slow-labelling but non-photosynthetic pool (2) would also be included in the estimate. The malate labelling kinetics in *S. viridis*, *P. miliaceum*, and *M. maximus* are also consistent with a more complex scenario with multiple pools of malate that all become labelled during a 60-min. pulse. We suggest that the new approach, comparing the initial labelling kinetics of the malate and aspartate pools, provides a more reliable estimate of the photosynthetically active malate pool size for flux estimation. The advantage is that it excludes metabolically active but non-photosynthetic pools of malate that equilibrate slowly with the photosynthetic pool from the calculations.

Using this approach, we observed large differences in the photosynthetically active malate pools, expressed as a percentage of total malate, in the two NADP-ME species: *Z. mays* (10%) and *S. viridis* (50%) (Figure 2B). The NAD-ME species, *P. miliaceum* (8%), had a similar relative pool size to *Z. mays*, while the PEPCK species, *M. maximus* (20%) had a somewhat larger photosynthetically active malate pool. The large pools of malate that are not directly involved in photosynthetic CO_2_ fixation complicate analysis of the malate labelling data to estimate photosynthetic fluxes. Therefore, we applied a correction to all of the malate labelling data using the estimated photosynthetically active pool sizes in the respective species.

To calculate the size of the metabolically inactive malate pool in absolute terms (i.e. nmol g^-1^FW), the amounts of unlabelled (m_0_) and labelled (m_1_-m_4_) malate isotopologues were summed and multiplied by the size of the inactive pool (expressed as a % of the total) derived from the linear regression analysis (Figure 2B). By this calculation, the photosynthetically active pools of malate in *Z. mays*, *S. viridis*, *P. miliaceum* and *M. maximus* were about 900, 1900, 280 and 500 nmol g^-1^ FW, respectively. In each species, the respective amount was then subtracted from the measured amount of unlabelled (m_0_) malate to give a corrected value for the amount of the photosynthetically active malate pool that was unlabelled (the calculations are presented in Supplementary Dataset S2A). After this correction, the abundance of the malate m_0_ isotopologue in the active pool declined in a monophasic manner during the ^13^CO_2_ pulse, falling to a near-zero value by about 120 s in all four species (Figure 2C). In parallel, the abundance of the malate m_1_ isotopologue rose rapidly to a peak representing 75-90% of the total photosynthetically active malate pool in each species, along with some label beginning to appear in the m_2_, m_3_ and m_4_ isotopologues (Figure 2C). This is consistent with the photosynthetically active malate pool rapidly becoming almost fully labelled in the C4 position. The labelling kinetics of the photosynthetically active malate pool closely resembled those of aspartate (Figures 2A and C).

The corrected m_0_ amount, representing the unlabelled portion of the photosynthetically active pool, was combined with the labelled isotopologue amounts to estimate total photosynthetically active malate pool sizes, isotopologue distributions and ^13^C enrichments (Supplementary Datasets S2B-D). The correction was applied to all samples from the pulse and pulse-chase labelling experiments. To calculate the average pool sizes of active malate in *S. viridis*, *P. miliaceum* and *M. maximus*, the amounts in all samples from the pulse and pulse-chase experiments were averaged for each species. As the *Z. mays* pulse (Medeiros *et al*., 2022) and pulse-chase (Figure 3) data were derived from separate batches of plants, we calculated the average photosynthetically active malate pool sizes separately for the two experiments on this species.

There were significant differences between species in the photosynthetically active malate pool size, as well as substantial variation within species, including a significant difference in *Z. mays* between the pulse and pulse-chase experiments (Figure 4). Although the *Z. mays* labelling experiments were performed at different times on two separate batches of plants, the plants were grown in the same growth chamber and, as far as possible, under identical conditions (pot size, irradiance, photoperiod, temperature, relative humidity). The composition of the soil and fertilizer were nominally the same in each experiment, but the plants were grown in different batches of soil. Therefore, it is possible that nutrient availability varied between the experiments, with differences in nitrate uptake, transport and storage potentially affecting malate pool sizes (Stitt *et al*., 2002). The photosynthetically active malate pool sizes were significantly higher in *Z. mays* (pulse-labelling experiment) and *S. viridis* than in *P. miliaceum* and *M. maximus*, but the latter two were not significantly different from the *Z. mays* pulse-chase samples (Figure 4). Therefore, despite a tendency for the NADP-ME species to have a larger photosynthetically active malate pool, we cannot make any firm conclusion that this is a significant difference from the other two subtypes. The aspartate pool size tended to be smaller in *Z. mays* than in *S. viridis*, *P. miliaceum* and *M. maximus* (Figure 4). The photosynthetically active malate pool was comparable to the aspartate pool in *Z .mays* and *S. viridis*, but was only about 20% of the size of the aspartate pool in *P. miliaceum* and *M. maximus*.

**Figure 4.**
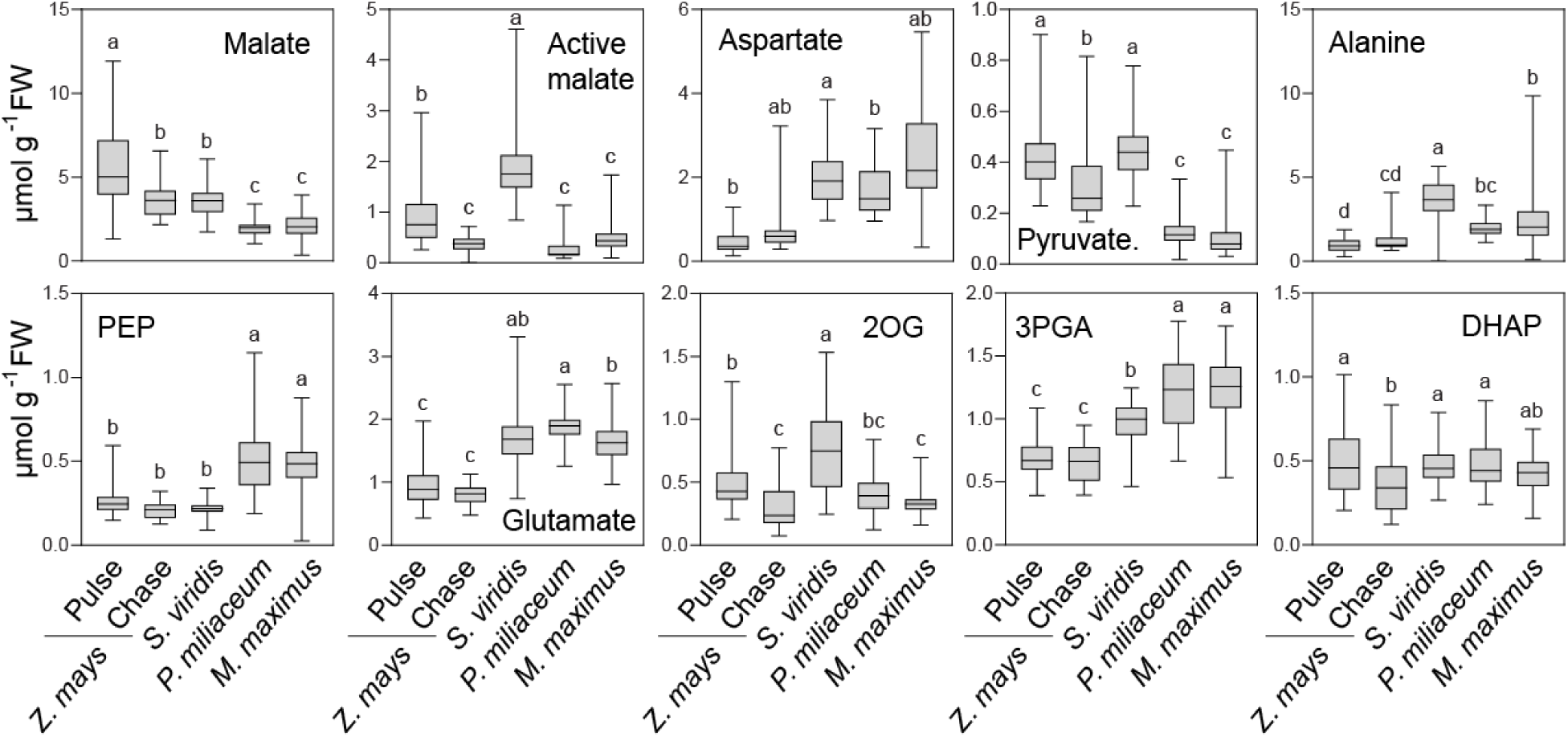
Comparison of metabolite pool sizes in C_4_ species representing different subtypes. The total pools of photosynthetic intermediates and associated metabolites were measured in leaves of *Zea mays, Setaria viridis* (NADP-ME subtype), *Panicum miliaceum* (NAD-ME subtype) and *Megathyrsus maximus* (PEPCK subtype) leaves from the pulse and pulse-chase labelling experiments presented in Figures 2-3. The data for *Z. mays* pulse-labelled samples are from a previous study (Medeiros *et al*., 2022). PEP, pyruvate and 3PGA pool sizes were measured enzymatically. For the other metabolites, the pool size was calculated by summing the measured amounts of unlabelled (m_0_) and ^13^C-labelled isotopologues. The photosynthetically active malate pool size was determined as described in Figure 2B. The data are presented as box plots (*n*= 21 to 66) showing the first quartile, median and third quartile, with the whiskers showing the minimum and maximum values. Significant differences between species according to one-way ANOVA followed by Tukeýs multiple comparison tests are indicated by letters. The original data are in Supplementary Dataset S2B.

### Labelling of PEP, 3PGA and other photosynthetic intermediates

With longer pulse times we observed transient rises in the abundance of m_2_ and m_3_ isotopologues of malate and aspartate, and a sustained rise in abundance of the fully-labelled m_4_ isotopologues (Figure 2A and C). This indicated gradual movement of ^13^C into the C1, C2 and C3 positions of the C_4_ acids. Carbons C1-C3 of malate and aspartate are derived from the PEP substrate of PEPC. In all four species we observed progressive labelling of PEP during the pulse (Figure 2D), accounting for the parallel appearance of label in the C1-C3 positions of malate and aspartate. During decarboxylation of the C_4_ acids in the BSCs, C4 is released as CO_2_, leaving a residual C_3_ moiety (C1-C3) in the form of pyruvate, alanine or PEP, which moves back to the MCs to provide PEP for further carboxylation. We observed progressive labelling of pyruvate and alanine during the pulse in all four species (Supplementary Figure S3), with similar kinetics to the labelling of PEP (Figure 2D). Together, these results show progressive labelling of the C1-C3 backbone of the C_4_ acids, and are consistent with rapid recycling of the residual C_3_ moiety from decarboxylation into PEP. The labelling of PEP showed similar kinetics to the labelling of 3PGA, but with a lag (Figure 2D).

During the pulse, the abundance of the unlabelled (m_0_) isotopologues of 3PGA, PEP, pyruvate and alanine declined towards zero in a monophasic manner, indicating that there were negligible inactive pools of these intermediates (Figure 2D and Supplementary Figure S3). Likewise, the pulse-labelling kinetics of other metabolites associated with photosynthetic metabolism – DHAP, glutamate and 2OG – did not show any evidence of inactive pools (Supplementary Figure S3). Therefore, unlike malate, there was no need to apply a correction before calculation of metabolically active pool sizes. The total amounts of pyruvate, PEP and 3PGA were determined enzymatically. Comparison of the *Z. mays* samples from the pulse and pulse-chase experiments showed no significant differences between experiments in the pool sizes of aspartate, alanine, PEP, glutamate and 3PGA, but there were significant differences for pyruvate, 2OG and DHAP (Figure 4). The pool sizes of PEP and 3PGA were consistently and significantly lower in the NADP-ME species compared to the NAD-ME and PEPCK species, while the pools of pyruvate were consistently higher in the NADP-ME species (Figure 4). Otherwise, the pool sizes generally overlapped across the subtypes.

### Estimation of C_4_ cycle carboxylation and decarboxylation fluxes

Photosynthetic metabolism is a dynamic process and quantifying fluxes is a key step in understanding how this metabolic network operates. Defining the active pool sizes of C_4_ pathway intermediates enabled us to estimate two key fluxes in the C_4_ pathway: flux of carbon into C_4_ acids via PEP carboxylation, and flux of carbon out of C_4_ acids during decarboxylation. To estimate the fluxes of carbon into malate and aspartate (“flux in”) during the initial carboxylation phase, we analysed changes in the levels of the m_1_ isotopologues of malate (corrected for the photosynthetically active pool size) and aspartate during the first 5 s of the pulse. As previously mentioned, malate and aspartate are expected to be labelled only in the C4 position during short ^13^CO_2_ pulses, so the m_1_ isotopologue data can be used as a reasonable proxy for specific labelling of malate and aspartate in the C4 position. We calculated the amounts of ^13^C in the C4 position of malate and aspartate at each time point by combining the m_1_ isotopologue abundance for each individual replicate with the size of the photosynthetically active malate and aspartate pools to estimate the amount of the m_1_ isotopologue. These were plotted against time and linear regression analysis was performed, fitting all of the data to a single line, to calculate initial rates for the first 5 s of the pulse, as described in Medeiros *et al*. (2022), with standard errors based on the 95% confidence interval. The data and calculations are presented in Supplementary Dataset S2F and Supplementary Figure S5A. To estimate the rates of release of ^13^C from the C4 positions of malate and aspartate during decarboxylation (“flux out”), we first calculated the amount of isotopologue in each replicate and at each time point from the start of the chase up to 120 s, and plotted them against time (Supplementary Figure S5B). We then fitted an exponential decay curve to the chase data for each species, and used the coefficients of the fitted curve to calculate the initial rate of C_4_ acid decarboxylation at time zero, i.e. the start of the chase, with standard errors based on the 95% confidence interval. The estimated fluxes into and out of the photosynthetically active malate and aspartate pools are presented in Figure 5 and Supplementary Dataset S2F, with separate calculations from the pulse (Medeiros *et al*., 2022) and pulse-chase experiments for *Z. mays*.

**Figure 5.**
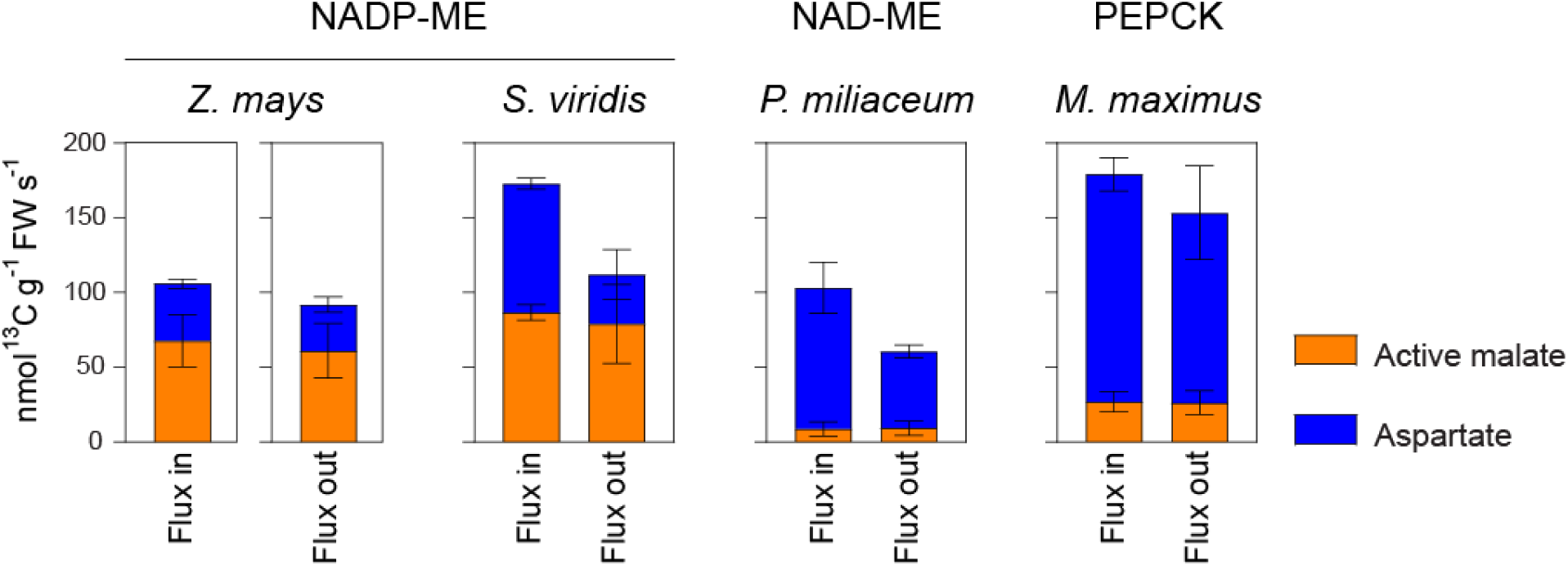
Estimation of carboxylation and decarboxylation fluxes in C_4_ species. The fluxes of carbon into and out of the photosynthetically active pools of malate and aspartate were estimated for *Zea mays, Setaria viridis* (NADP-ME subtype), *Panicum miliaceum* (NAD-ME subtype) and *Megathyrsus maximus* (PEPCK subtype). Estimates of fluxes into C_4_ acids (“flux in”) were calculated from changes in the levels of the respective m_1_ isotopologues during ^13^CO_2_ pulse labelling (0-5 s; Figure 2), serving as a proxy for specific labelling in the C4 position. “Flux in” values for *Z. mays* are derived from pulse-labelling data published in Medeiros *et al*. (2022). Estimates of fluxes out of C_4_ acids (“flux out”) were calculated from changes in the levels of the m_1_ isotopologues during the ^12^CO_2_ chase, after pulse-labelling with ^13^CO_2_ for 20 s (Figure 3A). For each species, a single exponential decay curve was fitted to the chase data for each metabolite (from 0-120 s), to calculate the initial rate of decarboxylation at the beginning of the chase (Supplementary Figure S5). The error bars represent the standard error of the mean based on the 95% confidence interval. For each species, the malate data were normalized to the respective photosynthetically active pool size (Figure 2B) prior to analysis. The original data are in Supplementary Dataset S2F.

In *Z. mays*, our estimates of flux into and out of C_4_ acids are in close agreement, with about two-thirds of the total flux being carried by malate and one-third by aspartate (Figure 5). In the other NADP-ME species, *S. viridis*, the flux of carbon into aspartate was greater, and approximately equal to the flux into malate (Figure 5). Our estimate of the flux out of malate in *S. viridis* matched that for flux into malate in this species, but the value we obtained for flux out of aspartate was lower than the estimated flux into aspartate. In *P. miliaceum* (NAD-ME), the vast majority of the C_4_ pathway flux was carried by aspartate, with only about 10% of CO_2_ being carried by malate (Figure 5). As for *S. viridis*, our estimate for the flux of carbon out of aspartate was lower than that for the flux in. Potential explanations for these discrepancies are discussed below. In *M. maximus*, aspartate was the dominant C_4_ acid, accounting for about 90% of carbon flux into and out of the C_4_ pathway while malate accounted for about 10% of these fluxes (Figure 5).

## Discussion

### Photosynthetically active pool sizes of C_4_ acids in C_4_ plants

The aim of this work was to explore the diversity of photosynthetic metabolism in C_4_ plants by measuring C_4_ pathway fluxes in representative species of the three C_4_ subtypes: NADP-ME (*Z. mays* and *S. viridis*), NAD-ME (*P. miliaceum*) and PEPCK (*M. maximus*). We applied classical pulse and pulse-chase labelling approaches to investigate metabolic fluxes, but using stable-isotope-labelled CO_2_ (^13^CO_2_) instead of the radioactive ^14^CO_2_ that was used in the experiments that led to the discovery and elucidation of C_4_ photosynthesis (Hatch and Slack, 1966; Johnson and Hatch, 1969).

We observed incorporation of label from ^13^CO_2_ into both malate and aspartate in all four species during the pulse-labelling experiments. The abundance of the unlabelled (m_0_) isotopologue of aspartate declined to near zero in a monophasic manner during pulse labelling up to 60 min (Figure 2A), indicating that essentially all of the aspartate in the leaf was present in the photosynthetically active pool. In contrast, a substantial proportion of the total malate remained unlabelled even after the leaves had been supplied with ^13^CO_2_ for 60 min (Figure 2A). This indicated that only part, indeed sometimes only a minor proportion of the total malate in the leaf was in the photosynthetically active pool. Therefore, to make any meaningful estimations of C_4_ pathway fluxes, it was essential to determine the sizes of the photosynthetically active malate pool in each species. To do this, we assumed that the initial labelling kinetics of the photosynthetically active malate pool would be similar to those of aspartate, given that both are derived from a common pool of OAA that is produced by the initial carboxylation reaction by PEPC (Figure 1A). We plotted the abundance of the m_0_ isotopologue of malate in each sample against the corresponding abundance of the m_0_ isotopologue of aspartate, and then used linear regression to fit a line to the data from the short-duration pulses (0-20s) (Figure 2B). The y-axis intercept of this line represents the time when all of the aspartate pool is at least partially labelled (i.e. every molecule contains at least one ^13^C atom). Given the common origin of the labelled aspartate and malate from labelled OAA, this also provides us with an estimate of the size of the corresponding malate pool that is actively involved in the C_4_ pathway.

In *Z. mays* (10%), *P. miliaceum* (8%) and *M. maximus* (20%), a minority of the total malate was in the photosynthetically active pool, whereas the active pool in *S. viridis* represented 50% of the total malate (Figure 2B). It is likely that most of the photosynthetically inactive malate is located in the vacuoles (Martinoia and Rentsch, 1994: Weiner and Heldt, 1987). When we compared the absolute sizes of the active malate pools (nmol g^-1^ FW), the NADP-ME species showed a tendency to have larger active pools of malate than the other two species, but the results were inconclusive due to differences between the two sets of *Z. mays* values (Figure 4). We also compared the sizes of the total malate pool and the photosynthetically active malate pool in the C_4_ species with the malate pools in a diverse range of C_3_ plants (Gerhardt and Heldt, 1984; Szecowka *et al*. 2013; Arrivault *et al*., 2015), where malate is not directly involved in photosynthetic CO_2_ fixation. Despite the very high flux through the photosynthetically active pool of malate in the C_4_ species, their active pools were noticeably smaller than the total pools of malate in the C_3_ species and the total pools of C_4_ species were in the same broad range as the total pools of C_3_ species (Supplementary Table S1), illustrating a general observation that pool size is an unreliable indicator of metabolic fluxes (Szecowka *et al*., 2013). Almost all of the malate is located in the vacuole in C_3_ species like spinach and Arabidopsis (Gerhardt and Heldt 1984, Szecowka et al., 2013) and up to 90% is in the vacuole in *Z. mays* (Weiner and Heldt, 1992).

### C_4_ pathway fluxes in NADP-ME, NAD-ME and PEPCK species

By normalizing the malate labelling data to the photosynthetically active pool size, we were able to estimate the metabolic fluxes into and out of malate in a way that allowed a meaningful comparison with the corresponding aspartate fluxes. In *Z. mays*, we found that the flux of carbon into C_4_ acids was mainly into malate (Figures 2 and 5). Nevertheless, there was a non-negligible flux of ^13^C into aspartate as well. Chapman and Hatch (1981) also reported substantial labelling of aspartate during ^14^CO_2_ pulse-labelling of *Z. mays* leaves. The malate dehydrogenase (MDH) and aspartate aminotransferase (Asp-AT) reactions are readily reversible, depending on the relative concentrations of the substrates and products, so there is likely to be some equilibration between the malate and aspartate pools, via OAA. Thus, it is possible for the aspartate pool to become rapidly labelled even if it is not carrying net flux through the C_4_ pathway. However, in the pulse-chase experiments we not only observed flux of ^13^C out of the aspartate pool during the chase but also noted that this was very rapid, being identical to or even marginally faster than that of malate (Figure 3A). This would not be expected if the aspartate were simply a dead-end pool that only becomes labelled via equilibration with malate in the MCs. This rapid loss of label from aspartate during the chase points to rapid movement of labelled aspartate from MCs to BSCs and decarboxylation of aspartate. Furthermore, the apparent rate of aspartate decarboxylation closely matched the flux of carbon into aspartate by carboxylation (Figure 5). It has previously been reported that there is an asymmetric distribution of aspartate in *Z. mays* leaves, giving rise to a concentration gradient that could drive diffusional movement of aspartate from the MCs to the BSCs (Arrivault *et al*., 2017). Together, these findings show that malate carries the majority (60-70%) of the C_4_ pathway flux in *Z. mays* leaves, as expected for an NADP-ME-type species (Figure 1A), but that there is also a substantial flux of aspartate through the C_4_ pathway in *Z. mays*, raising the question of how aspartate is decarboxylated in *Z. mays* leaves.

Chapman and Hatch (1981) reported that isolated *Z. mays* BSC strands could metabolize aspartate and 2-oxoglutarate to provide CO_2_ for photosynthesis. Wingler *et al*. (1999) confirmed this observation and showed that aspartate decarboxylation is inhibited by 3-mercaptopicolinic acid, a specific inhibitor of PEPCK. Hatch *et al*. (1975) found little or no PEPCK activity in leaves of *Z. mays* plants grown under natural high-light conditions, but other studies have reported readily detectable PEPCK activities in *Z. mays* leaves (Walker *et al*., 1997; 2002; Chao *et al*., 2014; Weissmann *et al*. 2016) suggesting that PEPCK could contribute to C_4_ acid decarboxylation in NADP-ME species (Pick *et al*., 2011). Chapman and Hatch (1981) proposed an alternative pathway for aspartate decarboxylation in *Z. mays*, with aspartate being deaminated by Asp-AT to OAA, which is reduced to malate by either the chloroplastic NADP-MDH or mitochondrial NAD-MDH, before decarboxylation of the malate by NADP-ME in the chloroplasts. In *Flaveria bidentis*, an NADP-ME-type eudicot, up to half of the C_4_ pathway flux is carried by aspartate and NADP-ME could also contribute to aspartate decarboxylation in this species (Meister *et al*., 1996). Whether PEPCK or NADP-ME mediates aspartate decarboxylation in *Z. mays* BSCs is uncertain, but has implications for the energetic cost of running the C_4_ cycle and redox homeostasis within the BSCs (Furbank, 2011). Reverse genetic approaches to selectively knock out enzymes or transporters needed by one or the other pathway would be one way to address this question, but such experiments are not trivial in *Z. mays* due to the difficulty of transformation.

In *S. viridis*, flux of carbon into aspartate represented about 50% of total carboxylation (Figure 5). However, our estimate of the fluxes out of C_4_ acids indicated that the rate of aspartate decarboxylation was only about half that of malate (Figure 5). Total decarboxylation flux was also only about two-thirds of the estimated carboxylation flux, but these should be equal if the plants were at steady state. This apparent discrepancy has been explored in depth in a parallel study (Baccolini *et al*., manuscript in preparation). In brief, we conclude that our calculations of decarboxylation flux can under-estimate the true decarboxylation flux due to leakage of ^13^CO_2_ from the BSC (Hatch *et al*., 1995) and re-fixation into C_4_ acids during the chase. Furthermore, there is a small amount of the m_1_ isotopologue of PEP (approx. 5% of the total PEP) present after a 20-s pulse with ^13^CO_2_ (Figure 2D), and carboxylation of this with ^12^CO_2_ during the chase will produce m_1_ isotopologues of malate and aspartate. By replenishing the pools of m_1_ malate and m_1_ aspartate during the chase, this will at least partially mask the loss of ^13^C from the original pools that were present at the start of the chase. It is not clear why these confounding effects appear to have a greater impact on our estimates of decarboxylation fluxes in *S. viridis* than in *Z. mays*. These complications should not affect our estimates of carboxylation fluxes, so we consider these to be the more robust indicator of the relative C_4_ pathway fluxes, and conclude that malate and aspartate fluxes are approximately equal in *S. viridis*. It has been reported that PEPCK transcript abundance is very low in *S. viridis* leaves (John et al., 2014; de Oliveira Dal’Molin 2016) and immuno-blotting with anti-PEPCK antibodies failed to detect PEPCK protein (Calace et al 2021; Bellasio and Ermakova, 2021). This suggests that that PEPCK makes little or no contribution to aspartate decarboxylation in this species, but the actual pathway remains unknown. Given that *S. viridis* is more readily transformable than *Z. mays*, it would appear to be a better model for reverse genetic approaches to investigate the pathway of aspartate decarboxylation in NADP-ME-type species.

Our estimates of carboxylation and decarboxylation fluxes indicate that aspartate is the dominant C_4_ acid in *P. miliaceum* and *M. maximus* (Figure 5), in broad agreement with the textbook models (Figure 1A). However, there are measurable carboxylation and decarboxylation fluxes into and out of malate in both species. Hatch *et al*. (1975) reported the following decarboxylating enzyme activities in *P. miliaceum* leaves: NADP-ME, 0.4 µmol min^-1^ mg^-1^ chlorophyll; NAD-ME, 4.8 µmol min^-1^ mg^-1^ chlorophyll; PEPCK <0.2 µmol min^-1^ mg^-1^chlorophyll. Assuming a chlorophyll content of 1.5 mg g^-1^ FW (Supplementary Figure S2), these translate to 10, 120 and <5 nkat g^-1^FW, respectively. The activities of NADP-ME and NAD-ME would be sufficient to mediate the malate and aspartate decarboxylation fluxes, respectively, that we estimated for *P. miliaceum* (Figure 5). In *M. maximus* leaves, Hatch *et al*. (1975) reported the following decarboxylating activities: NADP-ME, 0.2 µmol min^-1^ mg^-1^ chlorophyll; NAD-ME, 0.5 µmol min^-1^ mg^-1^ chlorophyll; PEPCK 9.8 µmol min^-1^ mg^-1^ chlorophyll, which translate to 7, 17 and 327 nkat g^-1^FW, respectively, if we assume a chlorophyll content of 2.0 mg g^-1^ FW (Supplementary Figure S2). This level of PEPCK activity would be more than enough to mediate the estimated aspartate decarboxylation flux (Figure 5), but the NAD-ME activity alone would be barely sufficient to mediate the estimate malate decarboxylation flux (Figure 5), suggesting that NADP-ME might also make a contribution.

### Re-fixation of CO_2_ via Rubisco

The classical ^14^CO_2_ pulse-chase labelling studies of Hatch and colleagues provided the crucial evidence for a key feature of the C_4_ pathway – the decarboxylation of C_4_ acids and re-fixation of CO_2_ into 3PGA via Rubisco (Hatch and Slack, 1966; Johnson and Hatch, 1969; Hatch, 1971). The pulse-chase approach, but using ^13^CO_2_ instead of radioactive ^14^CO_2_, was used in an attempt to detect C_4_-acid decarboxylation and CO_2_ re-fixation in transgenic *Oryza sativa* plants that had been engineered to express a minimal set of C_4_ pathway enzymes (Ermakova *et al*., 2021). However, no evidence of decarboxylation or re-fixation was found. The transgenic *O. sativa* plants had only low levels of expression of the heterologous *Z. mays NADP-ME* gene and only marginally increased NADP-ME activity compared to wild-type plants, which could account for the failure to detect C_4_ acid decarboxylation (Ermakova *et al*., 2021). The high rates of endogenous C_3_ photosynthesis occurring in the background of the *O. sativa* plants were an added complication, due to direct fixation of ^13^C into 3PGA, via Rubisco, and rapid equilibration of label between 3PGA and PEP, as discussed in Ermakova *et al*. (2021). These inconclusive ^13^CO_2_ pulse-chase labelling results left an unanswered question of whether ^13^CO_2_ pulse-chase labelling has sufficient sensitivity to detect C_4_ acid decarboxylation and re-fixation, as no C_4_ species was tested as a positive control. Therefore, the results we obtained from ^13^CO_2_ pulse-chase labelling in C_4_ species are pertinent for future investigations of C_4_ pathway fluxes in C_4_-engineered C_3_ plants.

We observed rapid loss of label from the C_4_ positions of malate and aspartate during the chase (Figure 3A), alongside either a transient rise in abundance of the m_1_ isotopologue of 3PGA (in *Z. mays* and *M. maximus*) or a lag at the start of the chase before the abundance of 3PGA (m_1_) started to decline (in *S. viridis* and *P. miliaceum*) (Figure 3B). The response of 3PGA was confirmed by a transient rise in labelling of DHAP in all four species (Figure 3B). These results show that stable-isotope pulse-chase labelling with ^13^CO_2_ does have sufficient sensitivity to detect C_4_ acid decarboxylation and CO_2_ re-fixation via Rubisco in authentic C_4_ plants. Therefore, at least in principle, this approach should also be suitable for detecting C_4_ cycle fluxes in C_3_ plants that have been engineered to operate the more efficient C_4_ pathway of photosynthesis.

## Acknowledgements

We would like to thank Laise Rosado de Souza for help with the gas exchange measurements and Bartimäus Jurke for help with sample processing.

## Author contributions

C.B., St.A., H.I., M.S. and J.E.L. conceived and planned the experiments. C.B. grew *Z. mays*, *P. miliaceum* and *M. maximus* and performed labelling experiments with these species. St.A. and H.I. grew *S. viridis* and performed the labelling experiments with this species. C.B. performed gas exchange measurements. C.B. and St.A. extracted samples for the LC-MS/MS analyses, which were performed by St.A. and R.F. C.B and St.A. processed the LC-MS/MS data and performed the spectrophotometric measurements of selected metabolites, chlorophyll and protein. C.B. and L.P.deS. extracted samples for GC-MS analyses which were performed by Sa.A. C.B., L.P.deS. and St.A. processed the GC-MS data. C.B. and St.A. prepared the figures and drafted the manuscript, J.E.L. revised the manuscript and all authors critically reviewed the manuscript before submission.

## List of Supplementary material

**Supplementary Figure S1.** Photosynthetic CO_2_ assimilation rates in C_4_ species.

**Supplementary Figure S2**. Leaf protein and chlorophyll contents in C_4_ species.

**Supplementary Figure S3**. ^13^CO_2_ pulse-labelling of photosynthetic intermediates and associated metabolites in C_4_ species.

**Supplementary Figure S4**. ^13^CO_2_ pulse-chase labelling of photosynthetic intermediates in C_4_ species.

**Supplementary Figure S5**. Analysis of ^13^CO_2_ pulse and pulse-chase labelling data to estimate C_4_ pathway fluxes in C_4_ species.

**Supplementary Table S1**. Malate and aspartate content of leaves from C_3_ and C_4_ species.

**Supplementary Dataset S1.** (**A**) Light saturation curve and (**B**) chlorophyll and protein contents.

**Supplementary Dataset S2.** (**A**) Isotopologue amounts, (**B**) total amounts, (**C**) isotopologue abundances, (**D**) ^13^C enrichments, (**E**) ^13^C amounts, and (**F**) fluxes in and out of the C4 position of active malate and aspartate.

